# Fast calcium transients in neuronal spines driven by extreme statistics

**DOI:** 10.1101/290734

**Authors:** Kanishka Basnayake, Eduard Korkotian, David Holcman

## Abstract

Fast calcium transients (< 10*ms*) remain difficult to analyse in cellular microdomains, yet they can modulate key cellular events such as trafficking, local ATP production by ER-mitochondria complex or spontaneous activity in astrocytes. In dendritic spines receiving synaptic inputs, we show here that in the presence of a spine apparatus (SA), which is an extension of the smooth ER, a calcium-induced calcium release is triggered at the base of the spine by the fastest calcium ions arriving at a Ryanodine receptor. The mechanism relies on the asymmetric distributions of Ryanodine receptors and SERCA pumps that we predict using a computational model and further confirm experimentally. The present mechanism where the statistics of the fastest particles arriving at a small target followed by an amplification is likely to be generic in molecular transduction across cellular micro-compartments such as astrocytes, endfeets and protrusions.

## 1 Introduction

Extreme statistics describes the distribution of rare events such as the first ions to find a small target 1,2], which are difficult to detect experimentally in biological microdomains. However, indirect signatures can be derived from their statistical properties. We examine here the role of rare events associated to calcium dynamics in dendritic spines, that are local microdomains located on the dendrites of neuronal cells. Spines can form synaptic connections that transmit neural activity 3, 4]. Here we describe specifically how the fastest calcium ions define the time scale of calcium transduction when it is followed by an amplification step. Regulating this fast event has many consequences in the induction of plastic changes. Indeed, calcium increase can be restricted to the spine head isolated from the dendrite, enabling the induction of local synapse-specific calcium-dependent plasticity leading to AMPA receptor accumulations 5, 6]. A fast and localized amount of calcium ions is necessary to induce ATP production from mitochondria to supply the energy required to maintain homeostasis 7-9].

Spines are characterized by the diversity of their shapes, sizes, and the presence or absence of different structural components and organelles such as an endoplasmic reticulum. During synaptic plasticity, spine morphology 10–12] can change, leading to an increase/decrease of the head size 13] or an elongation/retraction of their neck. Neck elongation can further lead to electrical and biochemical isolation from their parent dendrites 14–20]. Spines can contain a smooth endoplasmic reticulum (ER), fragmented in a compartment called the spine apparatus (SA), that can regulate calcium ions by storing or releasing them 21,22] and modulate synaptic inputs 23,24]. The SA is monitored by the actin-associated protein synaptopodin (SP) that can modulate calcium kinetics 25–27]. After calcium ions enter into dendritic spines, they can bind to endogeneous buffers, get extruded by pumps into the extracellular medium or be pumped into the SA by the sarco/endoplasmic reticulum calcium-ATPase pumps (SERCA3). Calcium ions can also induce calcium release from internal SA stores through the ryanodine receptors (RyR) 22,28]. However, the specific calcium regulation by SA remains unclear due to the fast dynamics and the spine’s nanometer-scale organization. We recall that the time scale of calcium diffusion transient during long-term plasticity 29] is of the order of hundreds of milliseconds 3, 30] but not faster.

Ve show here that the mechanism involved in fast calcium transient (faster than tens of milliseconds) relies on a new mechanism associated to the extreme statistics of the fastest ions that we describe here. For that purpose, we develop a computational model for calcium ion dynamics and use stochastic simulations to interpret uncaging and fluorescent imaging. To simulate calcium dynamics in synapses and dendritic spines, there are two possible approaches: deterministic reaction-diffusion equations 31,32] or stochastic modeling 33–37] that we use here. With this approach, we obtain a new understanding of fast calcium transients. Indeed, after calcium ions are released inside the spine head using flash photolysis of caged calcium, the concentration increase at the dendrite is faster in spines containing a SA compared to those where it is absent. This is a paradox as the SA should obstruct the passage from the spine head to the dendrite and prevent calcium ions from diffusing. To address this paradox, we use stochastic simulations to show that after calcium released in the spine head, the fastest ions arriving at the base determine the timescale of calcium transient due to an amplification step. Furthermore, we find that the distribution of arrival times of the fastest ions depends on the initial number of calcium ions, which is a signature of extreme statistics and rare events. Finally, we suggest that molecular activation initiated by the fastest particles is a generic mechanism in molecular transduction that can occurs in cellular micro-compartments such as protrusions, astrocytic endfeets and more. This mechanism is likely to define the timescale of biochemical activation in nano-and micro-domains, when the source of diffusing particles and the binding targets are spatially separated.

## Methods

### Stochastic simulations of calcium ions in a dendritic spine

Ve model a dendritic spine with a spherical head connected to a narrow, cylindrical neck 35] (Fig. 1A). The SA is also modeled with a similar geometry with a neck and a head positioned inside the spine (Fig. 2G). Calcium ions are described as Brownian particles following the Smoluchowski limit of the Langevin equation 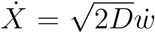, where *w* is the Wiener white noise. This motion of particles is simulated using the Euler’s scheme. Ions can diffuse inside the cytoplasm, and they are reflected at the surfaces of the spine and the SA (Snell-Descartes reflection).

**Figure 1:**
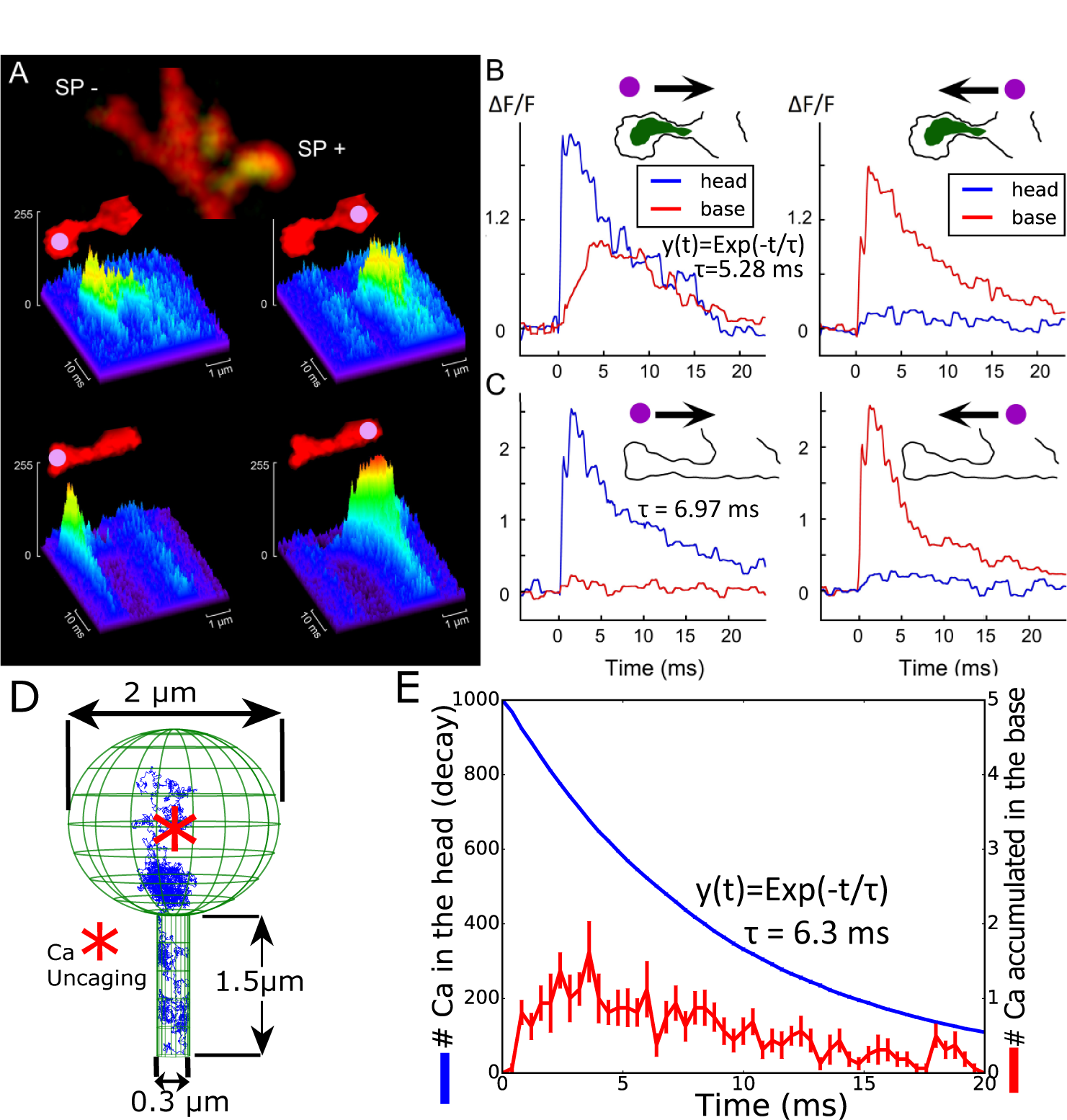
Calcium transient in dendritic spines with and without a SA. **(A)**. Examples of line scans in two neighboring spines of about same length, following flash photolysis of NP-EGTA in spine heads (left) and the parent dendrite (right). At the end of the experiment, the cultures were immuno-stained for SP (green staining). The same cell regions, containing recorded spines, were identified and imaged. Some spines (about 25%) of total mushroom spines contained synaptopodin puncta (SP+) while in the others (SP-) a clear puncta could not be seen. **(B) and (C)** Individual traces of calcium transients for the same spine heads (blue) and the parent dendrites (red) are shown on panel A. Schematic contours of two spines, containing (SP+) and not containing (SP-) synaptopodin puncta are shown on the top of the graphs. The arrows indicate the possible direction of calcium diffusion from the focus of uncaging (purple dots). There is a clear signal transfer from the spine head to the parent dendrite in the SP-positive spine and such signaling is absent if the focus of uncaging is set in the dendrite (B). For the SP-negative spine, calcium signaling in both directions looks the same (C). **(D)** Stochastic simulations of calcium ions in a dendritic spine without a SA: initial position of calcium ions (red star) and a trajectory (blue) are shown. The surface of the head contains 50 absorbing circular calcium pumps with a 10*nm* radius (not shown). **(E)** Simulated calcium transient following the model in (D). Calcium ions propagate from the head to the neck. The standard errors of the mean (5SEM) are shown by error bars.

**Figure 2:**
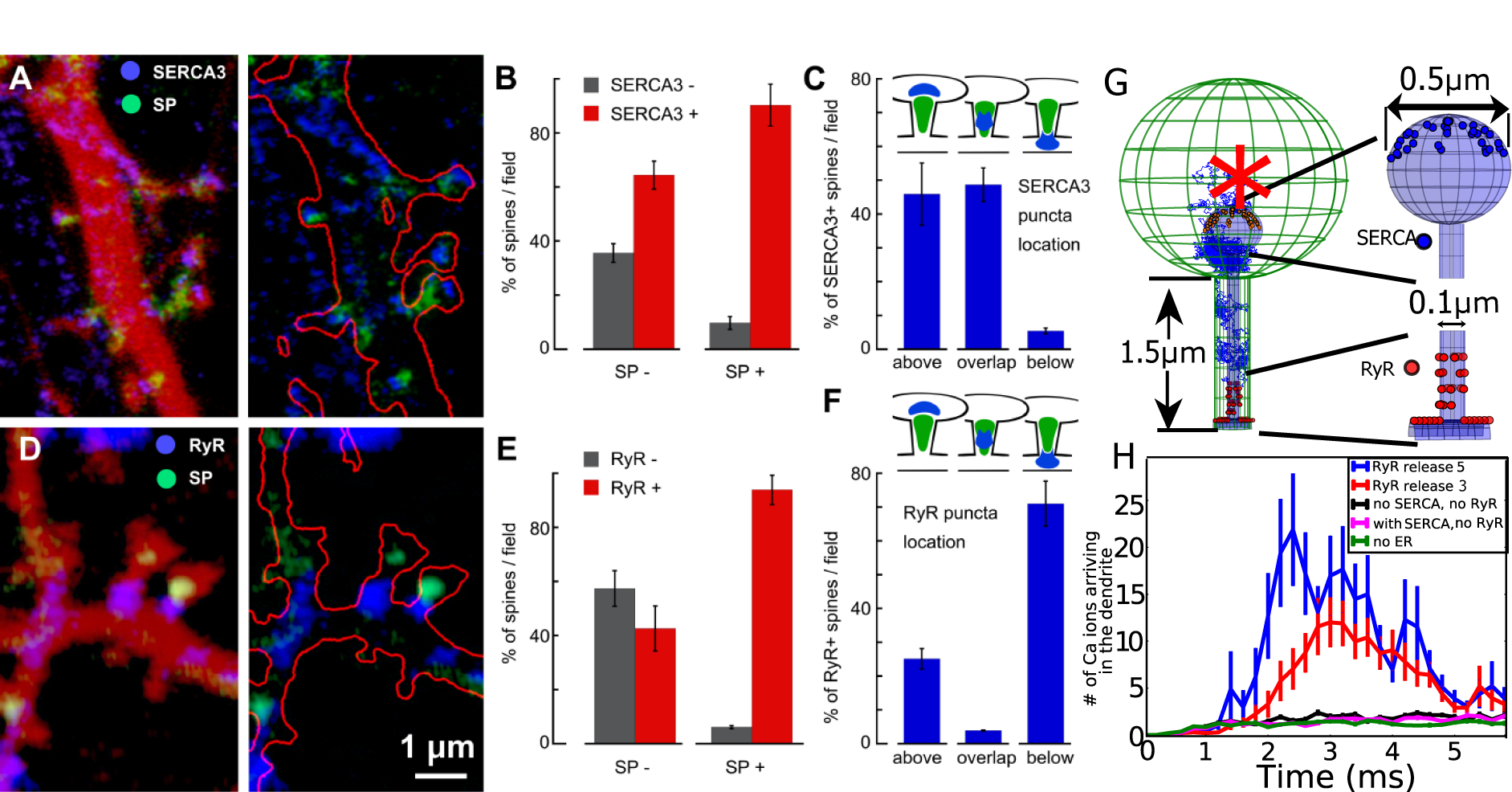
(A)-(F): Distributions of SERCA 3 and RyRs in dendritic spines depend on the presence of synaptopodin (SP) puncta. **(A)** and **(D)**-cultured rat hippocampal neurons were co-transfected with DsRed (red color on the right panels and red contour on the left panel) and SP (green puncta) and immunstained against SERCA3 and RyR. In SP occupied dendritic spines top head locations of SERCA3 (blue staining in (A)) and basal locations of RyR (blue staining in (D)) are confirmed. Gray columns versus red columns compares the presence of SERCA3 immunostaining **(B)** and RyR immunostaining **(E)** in SP-negative (left bars) and SP-positive (right bars) spines. Both the number of SERCA3-versus SERCA3+ as well as RyR-versus RyR+ spines is given in % per standard field (N-41 fields for (B) and 67 fields for (E)). Higher percentage of SERCA3+ and RyR+ spines in the SP+ groups (t-probability <0.01 for (B) and <0.001 for (E)). (C) Specific location of SERCA3 in SP+ dendritic spines. Number of spines is given as percentages of total SP+ and SERCA3+ spines per standard field, which is taken as 100%. Fields are the same as in (B). Schematic representations of SERCA3 and SP puncta aAIJtypicalaAi locations are shown on the top of the panel (C). Prevalence of SERCA3 puncta location above the SP puncta or overlapping with it (left and middle bars). **(F)** Location of RyR in SP+ dendritic spines. Again the number of spines is given as percentages when the total SP+ and RyR+ spines per standard field is taken as 100%. Fields are the same as in (E). Schematic representations are shown on the top of the panel (F). Prevalence of RyR puncta located below the SP puncta (right bar). **(G)-(H)** Stochastic simulations of calcium ions: (G) Model of a spine containing a SA with SERCA pumps and RyRs (right: magnifications show the distribution of 36 SERCA pumps in the upper hemisphere of the SA head and 36 RyRs located on the SA at the base, 12 on the shaft and 24 in the neck organized in four rings containing six randomly distributed receptors). Ions are initial released at the center of the head (red star). **(H)** Calcium ions arriving at the base of the dendrite following released in the head and released from RyRs (3 and 5 ions per activated receptors).

To replicate the uncaging experiments, we use either the center of the ball or the base of the cylindrical neck as the initial positions of particles. Ions arriving at the dendritic shaft located at the base are considered to be lost and do not return to the spine during the timescale of the simulations (absorbing boundary). The inner surface of the spine head contains 50 absorbing circular disks with a 10*nm* radius, which models calcium pumps.

### Ryanodine receptors

RyRs are activated upon the arrival of the first two ions 38] to a small absorbing disk of size *a*_*RyR*_ = 10*nm* (see SI for further descriptions). Ve positioned *n*_*R*_ = 36 receptors for the simulations organized in four rings in the SA neck, each containing six receptors. The other 12 are located on the SA component parallel to the dendritic shaft (see figure legends). After a receptor opens, it releases instantaneously a fixed number of calcium ions *n*_*Ca*_, which are positioned at the center of the receptors. Following this release, the RyR enters into a refractory period that lasts 3 to 6*ms*, during which it is modeled as a reflective boundary. In Fig 4B, we find that calcium ions should be released with a delay of 0.25ms after the arrival of a second ion to the RyR binding site.

**Figure 4:**
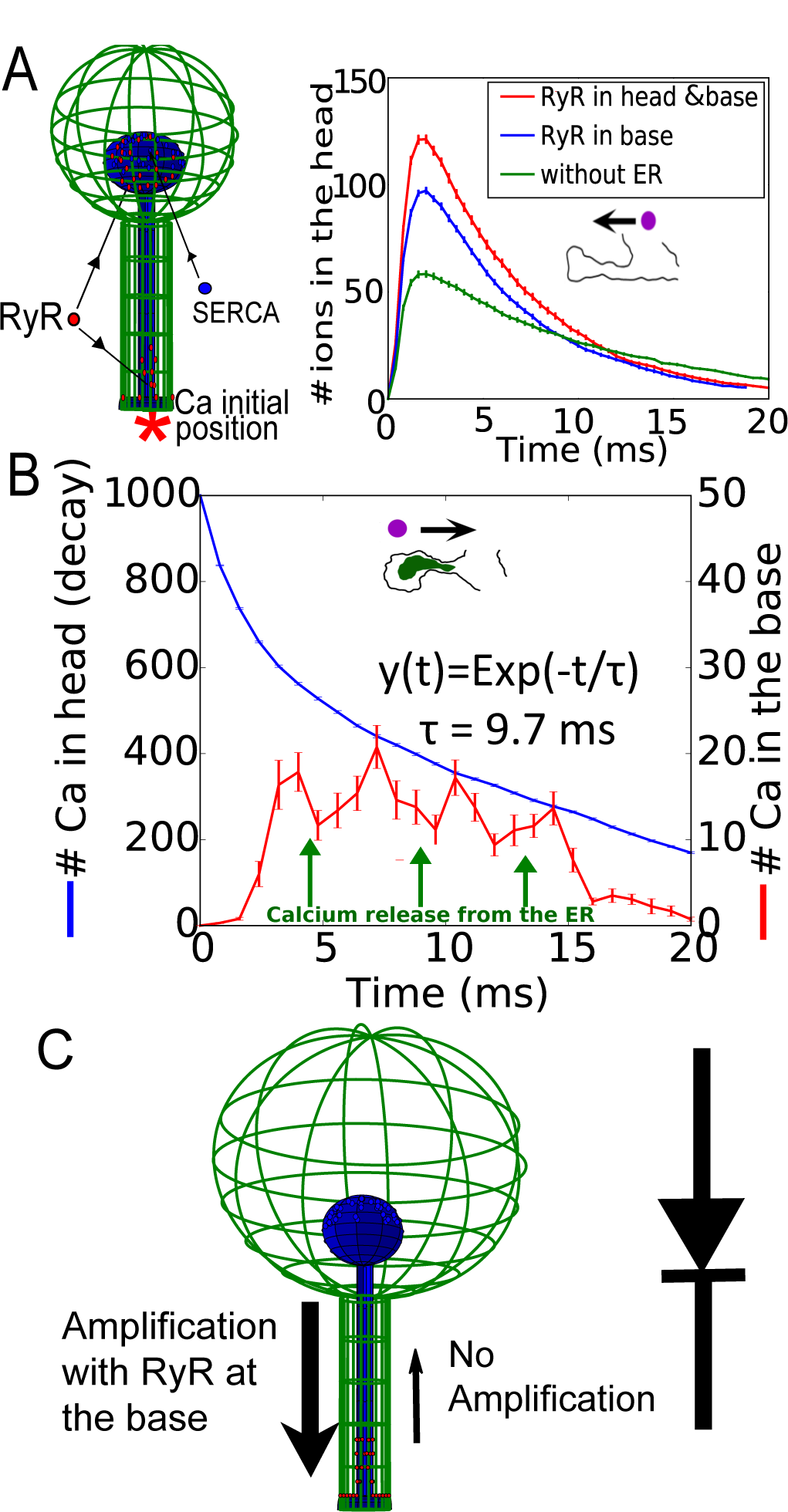
**(A) Left** Schematic representation of spine with SERCA pumps in the head and 36 RyRs placed in the head (18) and the neck (18). Calcium ions are released in the dendrite (red star). **(A) Right** Simulation of calcium transient in the head in the absence of a SA (green), when RyRs are present in both neck and head (red) and only in the neck (blue). **(B)** Stochastic simulations of calcium transient when taking into account SA depletion: RyRs are releasing with a delay of 0.25*ms*, initially 8, 7 and finally 6 ions with an averaged time indicated by the green arrows. The RyR refractory period is 3ms. **(C)** Diode representation of a spine with an SA.

### SERCA pumps

Ve positioned 36 SERCA pumps uniformly distributed on the upper hemisphere of the spine head. They are modeled as absorbing disks of size *a*_*SERCA*_ = 10*nm*. When a calcium ion arrives to the disk, it is bound indefinitely. If a second ion arrives, both are absorbed immediately and the transporter is frozen in an inactive state.

### Mean first passage time of ions to the base of a spine

For a Brownian particle released in the spine head, the mean arrival time to the base of a spine has been computed asymptotically 37]

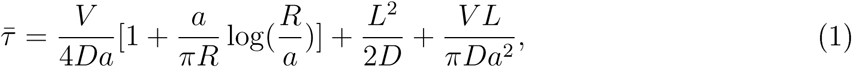

where *D* is the diffusion coefficient, a, R and L are the spine neck radius, head radius and the total length of the neck respectively. We refer to Table 1 (SI) for the parameter values, from which we estimated 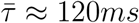.

### Experimental procedure

Cultured hippocampal neurons were transfected with DsRed and loaded with NP-EGTA AM (caged calcium buffer) after which several specific cells were microinjected with Fluo-4 calcium sensor. UV laser was directed to either spine heads or the basal dendritic shafts. Following the flashes of ND-YAG UV laser (4 ns, 330 nm) focused into a region of about 0.5 *µm* in diameter, the released calcium signals could be detected and line-scanned using confocal microscope at the rate of 0.7 ms/line. Other sources of calcium fluctuations were blocked using TTX (1 *µM*), DNQX (10 *µM*) and APV (20 *µM*). Following the experiment, cultures were fixed in 4% paraformaldehyde and immuno-stained for synaptopodin (green staining). The same cell regions, containing recorded spines, were identified and imaged.

## Results

### Fast calcium transient in spines with and without a SA cannot be due to classical diffusion

To investigate the role of the SA, we first released calcium following the flashes of ND-YAG UV laser to uncage calcium in dendritic spines from hippocampal neurons (Fig. 1A). After the experiment, the cultures were fixed using 4% paraformaldehyde and immuno-stained for SP to identify spines containing SA (see Material and Methods). Some spines (about 25%) of total mushroom spines contained synaptopodin puncta (SP +) while in the others (SP -) clear puncta could not be seen. Note that medium spines (of about 1-1.5 *µ*m in length) are studied. The transient fluorescence signal reveals the influence of the SA on calcium dynamics as shown in Fig. 1B-C. The calcium decay time in the head is well approximated by a single exponential 37] with a time constant *τ* = 5.28*ms* in *SP* + compared to *τ* = 6.97*ms* for *SP*, s<u>h</u>owing that the SA does not influence the extrusion rate from the spine head, probably because its obstruction is not completely occluding the passage from the head through the head-neck junction. However, the elevation of calcium at the base was much different, leading to high and very fast elevation in the case of SP+, a phenomena that is the focus on the present study. Finally, uncaging at the base of the spine leads to the same response in the head for SP+ and SP-, suggesting that the privileged calcium response occurs only in the head-neck direction when a SA is present. To further analyse the calcium transient, we use stochastic simulations, where ions are treated as Brownian particles (Material and Methods) and released at the center of the spine head (Fig. 1D-E). Using a single exponential approximation, we obtain a decay time in the head *τ* = 6.3*ms*. To conclude, fast calcium transient of less than 20*ms* in spines with no SP is well reproduced by stochastic simulations, but the calcium increase at the dendrite base for spines with a SA is much faster than the mean arrival time of calcium ions. This effect is surprising because it occurs despite the serious SA obstruction that should prevent calcium ions from passing easily through the neck. We shall now investigate the mechanism for this fast increase.

### The fast calcium transient is generated by calcium-induced calcium released and the asymmetric distribution of RyR on the SA

To clarify how the SA could affect the calcium transient, we first studied the distributions of SERCA3 and RyRs located on dendritic spines containing a synaptopodin puncta revealed by immunostaining Fig. 2A-D. We found that SERCA pumps are present in *SP* + spines and are located predominantly above the SA inside the spine head (Fig. 2A-C). This is in contract with the distribution of the RyRs, present in the *SP* + and mostly located below the SA at the base of the spine neck (Fig. 2D-F).

To study the consequences of these SERCA and RyR distributions on calcium transients, we run stochastic simulations similar to the ones we used in Fig. 1F. Ions are released initially in the spine head (red star Fig. 2G). 36 SERCA pumps (blue) are located on the surface of the SA (head) and 36 RyRs are located on the SA at the base (red). While SERCA pumps can uptake calcium ions from the cytoplasm to the ER, RyRs generate a calcium flux from the SA to the cytoplasm when two calcium ions are bound (Material and Methods). Interestingly, and in contrast to the results of Fig. 1E, after 1000 ions are released in the spine head, a significant calcium increase can be observed at the base of the spine in less than 2*ms*. This effect is already present when 3 calcium ions per RyR are released (Fig. 2H), and further amplified with 5 ions, compared to spines with no SA (green curve). At this stage, we conclude that calcium increase at the base of the spine is due to the calcium release from the SA. This release is induced by the opening of RyRs and triggered by the fastest calcium ions traveling from the head to the neck inside the cytoplasm.

### Extreme statistic for the fastest ions as a mechanism for activating Ryanodine receptors during calcium transients

To clarify the fast calcium transient observed at the base of a spine, we studied using modeling and simulations the dynamics of RyR opening when ions are released inside the head (Fig. 3A). In this stochastic model, a RyR opens when two ions are bound (Fig. 3B) (also Material and Methods) and releases calcium ions from SA to the cytoplasm Fig. 3B. Stochastic simulations reveal that in the presence of RyRs, the calcium released in the spine head induces a calcium increase at the base within the first 5*ms* (Fig. 3C). This effect is modulated with the distribution of SERCA pumps, but was clearly due to the presence of RyRs. This result confirms the role of RyRs in generating the fast calcium transient at the base of a spine.

**Figure 3:**
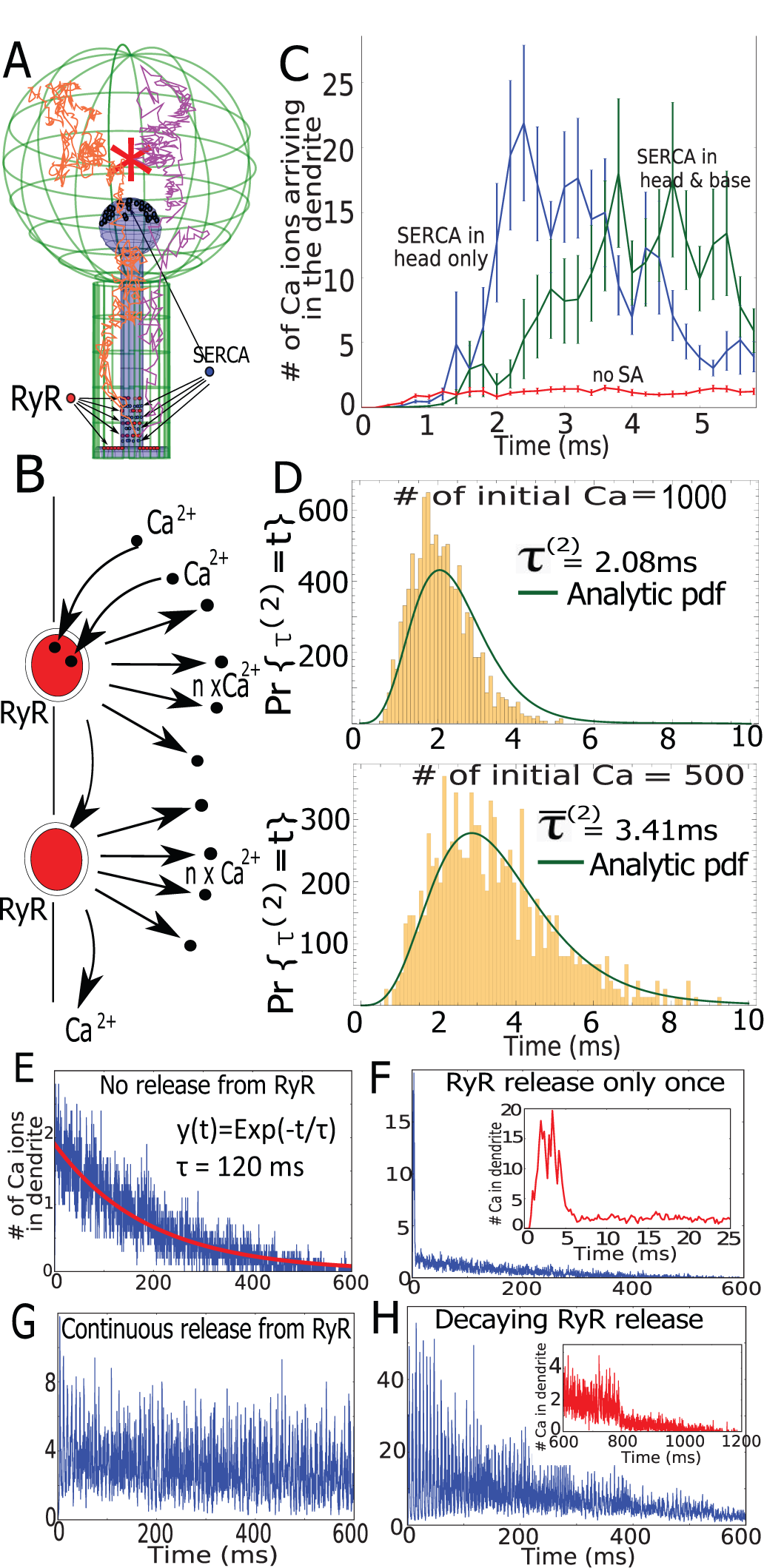
**(A)** Schematic representation of calcium trajectories in a spine with ER, where we placed SERCA (blue) in the head and at the base. They are arranged in four separated layers containing nine of them. RyRs arrangment is described in Fig. 1H. **(B)** Schematic release of calcium ions from RyR opening, triggered by the arrival of two calcium ions. **(C)** Dynamics of calcium accumulation in the dendrite in the presence (green) or absence (blue) of SERCA pumps at the base compared to the dynamics in a spine with no SA (red). **(D)** Histograms of arrival times of the two fastest calcium ions that open the first RyR when the initial numbers are either *N* = 1000 and *N* = 500, super-imposed with the analytical solution (not a fit) of Equation_8_ 5. **(E)** Transient of calcium ions arriving at the base of the dendrite when no RyRs are present, but no SERCA and calcium pumps. **(F)** Similar to (G) but all RyRs open and release Ca for only once and then deactivated (magnified in the inset). **(G)** Continuous release of two ions from each RyR with a refractory period of 6ms. **(H)** Same dynamics as in (G), but the number of releases from RyRs is reduced exponentially with time. Vhen a RyR becomes open, there are *n*_*Ca*_ ions released and the receptor is desensitized for 6*ms*. Initially *n*_*Ca*_ = 8 and decreases exponentially with a timescale of 360*ms*.

To assess the timescale of RyR activation, we constructed histograms of the first time *τ* ^(2)^ to activate RyRs by the binding of two successive calcium ions and the distribution is shown in Fig. 3D. Interestingly, the distribution depends on the initial released calcium concentration. We computed numerically from the histogram the mean opening times which is 3.4ms (resp 2.1ms) when 1000 (resp. 500) ions are released. These times are much faster than the mean time for an ion to arrive at the base of a spine in the order of 120*ms* (equation 1). We can now understand that this fast ion arrival is located in the tail of statistical distribution, which therefore selects the fastest among many Brownian particles.

### General theory of extreme statistics for Brownian calcium ions in a cellular microdomain

To further validate the results of stochastic simulations described in the previous section, we computed from the distribution of arrival time for *N* independent Brownian trajectories (ions) at a small binding site inside a bounded domain Ω. This time is defined by *τ* ^1^ = min(*t*_1_, …, *t*_*N*_), where *t*_*i*_ are the arrival times of the *N* ions. The arrival probability can be computed when the boundary *∂*Ω contains *N*_*R*_ binding sites *∂*Ω_*i*_ ⊂ *∂*Ω so that the total absorbing boundary is 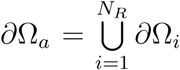, and the reflecting part is *∂*Ω_*r*_ = *∂*Ω − *∂*Ω_*a*_. The probability density function of a Brownian motion is the solution of

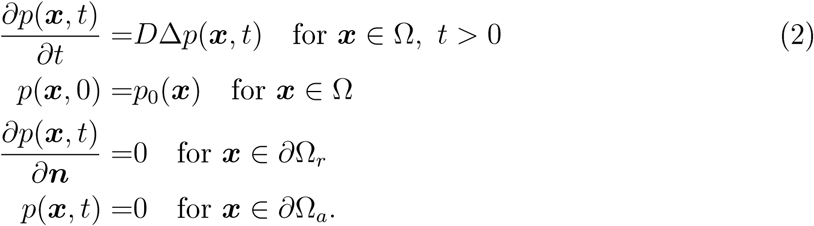

The survival probability, which is the probability that an ion is still not absorbed at time *t* is given by

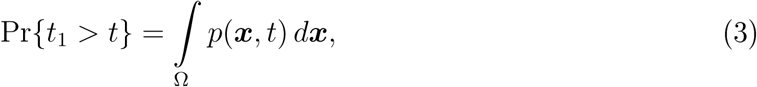

so that 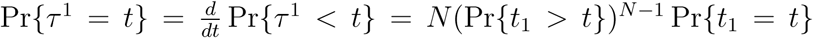, where 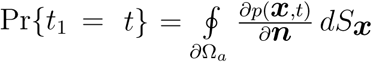 and 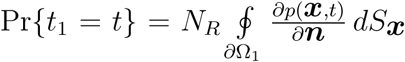. Putting all the above formula together, we obtain that the pdf for the first particle to arrive is

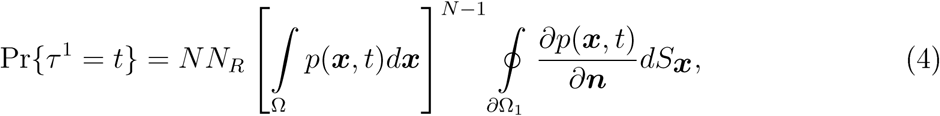

and the arrival time *τ* ^(2)^ for the second ion, which modeled the activation of a Ryr, is that of the minimum of the shortest arrival time in the ensemble of *N* − 1 trajectories after the first one has arrived and is given by

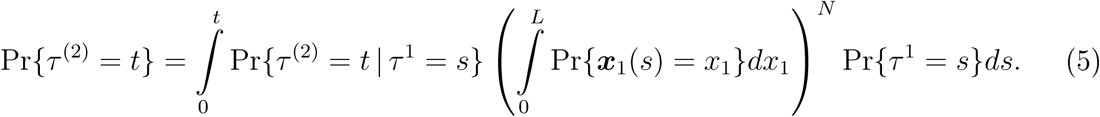

Ve plotted in Fig. 3D the solution of equation 5 where the distribution of arrival time for the first ion Pr {*τ* ^1^ = *s*} also accounts for the return of the ion located in the neck back to the head (see SI for the complete derivation). Ve find that the analytical solution superimposes with the stochastic simulations, confirming the consistency of the stochastic simulations and the theory of the extreme statistics. To conclude, this analytical approach further confirms the role of the fastest ions in setting the timescale of calcium-induced calcium released by RyR activation. We observe that the typical shortest path is very close to the shortest geodesic going from the initial position to the RyRs, which is much different compared to the paths associated with the mean arrival time.

### Long-time dynamics of calcium induced calcium-release

To further confirm the role of the fastest ions in triggering calcium release, we generated much longer simulations over 600ms (Fig. 3E-H). In the absence of an SA, we simulate the flux of ions arriving at the dendrite, showing an exponential decay of *τ* = 120*ms* (Fig. 3E), indeed in agreement with equation 1. Ve note that here there are no extrusion mechanisms such as calcium pumps. To evaluate the impact of the SA, we simulated a single release event of calcium ions, following RyR activation (5 ions are released per receptors) Fig. 3F. This release is local and affects only the global decay during the first few milliseconds (insets).

Vhen RyRs are releasing a minimum number of 2 calcium ions with a refractory period of 6*ms*, after a fast transient regime, the ensemble of RyR self-entertains (Fig. 3G). Indeed, when the SA contains a sufficiently large amount of calcium, the locally released calcium binds to RyR that opens, but the ions disappearing at the base of the spine are not sufficient to prevent this positive feedback loop between calcium and RyRs.

Finally, to account for a local SA calcium depletion, we simulated calcium release when it decreases in time starting from 8 ions with an exponential decay time of 360*ms*. After 600*ms*, the transient regime disappears (Fig. 3H).

To conclude, the two fastest ions arriving at a single RyR trigger the release of calcium from SA that induces a local calcium release. The timescale of activation depends on the initial number of released calcium ions in the head, which is the signature of an extreme statistics mechanism. This avalanche mechanism is responsible for the fast and large calcium increase at the base of the spine, when ions are diffusing from the head. Thus a release of local calcium ions from RyRs amplify the calcium signal. Furthermore, the calcium transient termination can be attributed to the local SA depletion over a few hundred milliseconds.

### Asymmetric calcium dynamics between spine and dendrite

To investigate the consequences of RyR distribution on calcium transients, we replicate the experimental protocol described in Fig. 1B-C with numerical simulations. We run simulations using the numerical scheme as the one described in Fig. 1D, with SERCA pumps located on head of the SA, while calcium ions are released at the base of the spine (red star in Fig. 4A). We tested two RyR distributions: (1) RyRs are only at the base of the neck, as suggested from Fig. 2F (2) RyRs are located also in the spine head.

Ve find that adding RyRs only at the base already increases significantly the calcium transient in the spine head (blue vs green, Fig. 4A). If in addition, RyRs are added in the head, the calcium transient in the head is further increased (red versus blue). However in this direction, since calcium transient in the head is not amplified as confirmed by our experimental findings (Fig. 1B-C), the present stochastic simulations agree with the immunostaining results of Fig. 2F, suggesting that there should be no RyRs in the spine head. We thus conclude that the asymmetric distributions of the RyRs contribute to the asymmetry of calcium transmission.

At this stage, we could not access to the calcium dynamics in the ER. However, it could have a drastic consequence on the calcium transient, as shown in Fig. 3E-H. We thus decided to use the transient fluorescence signal (Fig. 1B red curve) to recover the local SA depletion at a timescale of 20*ms* following calcium release. Releasing consecutively 8, 7 and then 6 ions per RyR with a refractory period of 3ms between each release, we could recover the transient kinetics observed experimentally (Fig. 4B red).

Ve also found here that calcium release from the RyRs is delayed by 0.25*ms* following the binding of two calcium ions. These results give us an indication of the SA depletion timescale, which is at a few tens of milliseconds. Putting the present results together, we find that the SA plays the role of a diode, amplifying ions from the head to the dendrite, but not in the opposite direction (Fig. 4C).

## Discussion

In the present study, we investigated how SA influences calcium transient inside dendritic spines. We found that calcium increases at the dendrite following uncaging in the head occurs in a few millisecond timescale. As shown here, this property can be explained by the statistics of the fastest calcium ions, without the need of considering electro-diffusion 19, 39, 40]. The fastest arriving ions open RyRs located at the base of the spine neck, which generate an avalanche through a calcium-induced calcium release from SA. This local avalanche leads to an accelerated amplification of the calcium signal before the remaining ions diffuse from the head.

Furthermore, the comparison of stochastic simulations with the experimental calcium transient constraints the number of ions released by the opening of a RyR: indeed we find that releasing 3 to 8 ions each time a RyR opens gives us a range of concentrations for experimental observations. Interestingly, we could recapitulate the decrease of released calcium by decreasing the released number from 8 to 6 one-by-one, occurring once in a few milliseconds (Fig. 4B). The distribution of RyRs has little influence on the calcium transient, as long as their inter-distances are less than few tens of nanometers (see also SI Fig. 3).

### Role of extreme statistics in molecular transduction in nanodomains

As RyRs are activated by the two fastest ions that arrive to the binding sites, the physical separation of the initial calcium release in the spine head and the location of the RyR in the base is compensated by the redundancy based on the large initial number of released ions. The timescale induced by the fastest ions depends on the logarithm of initial ion number and the length of the direct ray starting from the source and ending at the target 41].

The timescale generated by the fastest particle or ions is generic and can occur in many molecular transduction pathways where there is a separation between the initial source and a second step that consists of amplifying the signal. This is the case for second messengers such as IP3 42], G-protein coupled receptors 43] or modulation of the inner hair cell voltage by calcium-induced calcium release 44]. This time scale is very different from the Narrow Escape Time 37] phenomena, where the timescale depends on the volume of the domain. We conclude that the statistics of the fastest particle compensates for long distances and the key modulating parameter is the number of initial ions.

### Consequences of amplifying calcium concentration at the base of a dendritic spine

Vhat could be the role of calcium signal amplification induced by SA release at the base of a dendritic spine and not in the head? The asymmetry of RyR localization is a key feature in this difference, leaving the head compartment separated from the rest of the dendrite. Amplifying calcium at the base could favor receptor trafficking by influencing the delivery of AMPA receptor to dendritic spines. Successive calcium accumulative events leading to SA refilling could trigger a massive release, while a depleted SA would only lead to a small release, suggesting an integrating role of the SA. Experimental evidences 45] further suggest that gating of the RyR is also modulated by the luminal calcium concentration fluctuations while the release can diminish when calcium is bound to buffers such as calse-questrin 46]. Another consequence of amplifying locally the calcium concentration at the ER is to trigger the production of ATP from nearby mitochondria 8]. Indeed, inducing ATP production requires that the calcium concentration reaches a threshold of 10*µM*. We studied here a timescale of few to 20 milliseconds. For longer durations, other mechanisms such as secondary messenger involving IP3 receptors 42] or the ORAI1 pathways 47] involved in SA replenishment can contribute to calcium concentration regulation. Future models should also incorporate the cycle of SA calcium, depletion using the ORAI1 pathways and the calcium uptake at the base of the spine by mitochondria 48].

## Author Contributions

All three authors designed the study. KB and DH conducted the mathematical analysis and computational simulations. EK performed the experiments and analysed the data. KB and DH wrote the paper.

## Acknowledgements

This research was supported by the Fondation pour la Recherche Medicale - Equipes FRM 2016 grant DEQ20160334882.

